# Root exudation and rhizosphere microbial recruitment are influenced by novel plant trait diversity in carrot genotypes

**DOI:** 10.1101/2024.03.12.583384

**Authors:** Hannah M. Anderson, Grace A. Cagle, Erica L.-W. Majumder, Erin Silva, Julie Dawson, Philipp Simon, Zachary B. Freedman

## Abstract

Root exudate composition can influence rhizosphere microbial recruitment and is tightly controlled by plant genetics. However, little research has profiled root exudate in vegetable crops or determined their role in rhizosphere microbial community and metabolite composition. It is also not well understood how root exudates and resulting rhizosphere dynamics shift across plant trait diversity and with the development of novel crop genotypes. To address these knowledge gaps, this study paired metabolomics and microbiome analyses to evaluate associations between the composition of exudates, soil bacterial and fungal communities, and soil metabolites across four genotypes of organically produced carrot of differential breeding histories, including two experimental genotypes. Plant genotypes modified soil microbial diversity and composition, and differentially recruited bacterial taxa with demonstrated potential for plant-growth related functions including ammonia oxidation, nitrogen fixation, and phytohormone production. Bacterial rhizosphere recruitment from bulk soil was genotype and root exudate-mediated, while fungal recruitment was not. Moreover, root exudate composition was distinct in an heirloom genotype and a novel nematode resistant genotype, compared to other genotypes tested. Root exudate and rhizosphere metabolite composition was decoupled, and soil metabolites strongly associated with fungal, but not bacterial communities. Taken together, the results of this study suggest that novel crop trait diversity and breeding histories hold consequences for the functional potential of soils through the diversification of root exudate mediated plant-microbe interactions.

## 1. Introduction

Terrestrial plants evolved in close association with soil microorganisms, mediating both positive and negative plant-microbe interactions through root growth and rhizodeposition. The rhizosphere, defined as the narrow zone of soil in direct contact with an actively growing plant root, is a hotspot for plant-soil substrate flow and microbial activity (Kuzyakov & Blagodatskaya 2015). It is well documented that rhizosphere microbial community composition and functions differ from that of the bulk soil, and these differences could ultimately shape plant and soil functional outcomes as a result of shifts in microbial metabolism, for example, by increasing soil nutrient availability (Ling et al. 2022). While the bulk soil microbial community initially shapes the rhizosphere as the “microbial seed bank”, observed differences in rhizosphere microbial community composition between plant species and even genotypes during plant development indicate a strong host plant-specific selective pressure (de Ridder-Duine et al. 2005, Micallef et al. 2009, Aira et al. 2010). Oftentimes, the specific plant or soil-based mechanisms resulting in these observed patterns of rhizosphere microbial community assembly remain elusive.

The flux of photosynthetically fixed carbon (C) from plant tissues to the rhizosphere through root exudation is a potential mechanism that can modify soil microbial abundance and activity. As a source of low-molecular weight (LMW) labile C, root exudates provide a readily accessible energy source for microbial metabolism, which can ultimately stimulate microbial recruitment and growth (Meier et al. 2017, Zhou et al. 2022, Liu et al. 2022). Early research on plant root exudation provided the foundational understanding that root exudation is significantly controlled by plant genetics (Lynch & Whipps 1983, Larsen et al. 1998, Tadano et al. 1993, Kamilova et al. 2006). More recent applications of high-throughput metabolomics and microbial community analysis support the hypothesis that root exudate chemical identity regulates specific microbial selection in the rhizosphere through preferential uptake and metabolic use (Broeckling et al. 2008, Eilers et al. 2010, Hugoni et al. 2018, Zhalnina et al. 2018). Additionally, soil microorganisms produce and excrete LMW metabolites that can shape the biochemical habitat and carbon availability in the rhizosphere (Bi et al. 2020, Swenson et al. 2015, Song et al. 2020). Despite the apparent importance of both root exudates and microbial metabolites, there is a lack of research that has determined the influence of root exudates on soil metabolite composition or investigated both root and soil metabolomes in tandem to differentiate their effects on rhizosphere microbial communities. Metabolomics-based root exudate profiling to date has been mainly applied to grasses and crops used for bioenergy such as switchgrass and sorghum, and plants are usually grown hydroponically or in artificial soil (Miller et al. 2019, Dietz et al. 2020, Seitz et al. 2022). There is notably little application of these methods in vegetable crops, especially in field soil environments, significantly limiting the integration of developments in plant-microbe interactions into food system sustainability (Neumann et al. 2014, Zhao et al. 2023).

Cultivars developed by plant breeding efforts introduce phenotypic diversity into agricultural systems, potentially altering rhizosphere functioning through shifts in plant-microbe interactions across plant genotypes. For example, domestication can alter root exudate profiles and rhizosphere microbial interactions, suggesting an influence of breeding history in some crops (Iannucci et al. 2017, Yue et al. 2023). However, whether crop trait diversity affects rhizosphere plant-microbe interactions in vegetable production systems, and if novel trait introductions can modify these interactions, has not been determined. An increased understanding of these effects is important, as crop trait diversity can influence soil biological health by modifying microbial diversity, activity, and associated carbon and nutrient cycles (Tiemann et al. 2015, Wood et al. 2015, Cortois et al. 2016, Singh et al. 2018, Zhang et al. 2021, Koyama et al. 2022). These effects are likely to be especially important in agricultural systems with heightened reliance on soil microbial activity for agronomic outcomes, such as organically managed soils, which is a rapidly expanding land use (USDA 2021).

In organic agriculture, carrot (*Daucus carota* subsp. *sativus*) is one of the most highly produced vegetables in the United States, spurring the establishment of breeding programs prioritizing the development of cultivars that are well-adapted to organic growing conditions and meet market demand (USDA 2022, Simon et al. 2016, Simon et al. 2021). Pathogenic nematode resistance and enriched anthocyanin content associated with purple coloration are presently being explored in experimental carrot lines. These ongoing carrot breeding efforts have demonstrated potential impacts to rhizosphere functioning in climate-controlled field and greenhouse settings, making carrot a suitable candidate for exploring plant-microbe interactions across trait diversity in organic vegetable systems in a field setting (Keller-Pearson et al. 2020, Triviño et al. 2023). The mechanisms driving these observed shifts in plant-microbe interactions across carrot cultivar development are currently unknown. Early research using carrot as a model plant has suggested that root exudation mediates interactions with arbuscular mycorrhizal fungi, suggesting a potential role for these compounds in rhizosphere signaling (Bécard & Piché 1989, Nagahashi et al. 1999, Nagahashi et al. 2000, Nagahashi et al. 2007, Poulin et al. 1993). However, root exudates have never been fully profiled in carrots and the biochemical composition of root exudates in vegetables in general is severely understudied.

The aim of this study was to advance understanding of the mechanisms that shape rhizosphere microbial communities by co-investigating the diversity of root exudate composition, soil metabolomes and rhizosphere microbial recruitment in a field setting. In this way, the objectives of this study were to 1) assess root exudate and soil microbial communities across genotype diversity in organically produced carrot, and 2) evaluate whether root exudate or soil metabolites modulate soil microbial assembly in the rhizosphere. Related to objective 1, we hypothesized that 1.1) across genotypes, plant genetic differences will lead to distinct root exudate and soil microbial community composition and 1.2) rhizosphere microbial recruitment will shift with genotype across the growing season. Related to objective 2, we hypothesized that 2.1) root exudates will modulate soil metabolite composition and 2.2) differences in root exudate and soil metabolite composition across plant genotypes will account for variation in both bacterial and fungal rhizosphere communities.

## 2. Methods

### 2.1 Genotype Selection

To determine the effect of plant trait diversity on soil microbial community assembly, four genotypes of carrot were selected for a field trial based on previous research and current breeding efforts (Simon et al. 2016, Keller-Pearson et al. 2020, Simon et al. 2021, Triviño et al. 2023). The selected genotypes are phenotypically diverse in terms of root shape, flavor, and color, and include one heirloom variety (Red Core Chantenay, hereafter “H”), one hybrid variety (Bolero, hereafter “Hy”), an experimental nematode resistant line (Nb8503, hereafter “NR”), and an experimental purple/orange-cored carrot (P0114, hereafter “PO”) (Table 1).

**Table 1.**
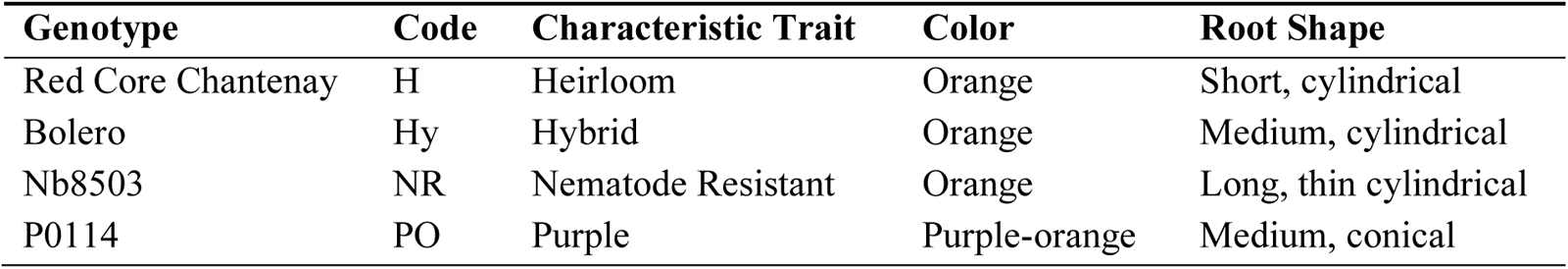
Assigned traits of each carrot genotype. Acquired from https://carrots.eorganic.info/.

### 2.2 Field Trial

A randomized complete block design field trial (n=5 blocks) was conducted at West Madison Agricultural Research Station on certified organic land. The soil is classified as a Kegonsa silt loam and has been cultivated under a diverse organic crop rotation. Crop rotation history and manure inputs for six years are available in Table S1. Soil chemistry data from annual soil tests including soil pH, organic matter (%), P (ppm) and K (ppm) is available in Table S3. The experimental plots received no external fertilizer amendments the year of planting. Prior to planting, the site was plowed to a depth of 15 cm and existing cover crop residue was incorporated into the soil. Carrots were established from 6/20/22 to 9/26/22 in 1m^2^ plots seeded at a rate of 100 seeds/meter using methods that are common in regional production systems. Drip irrigation was delivered daily. In total, 4 genotypes with 5 replicates yielded 20 experimental plots.

### 2.3 Midseason and Harvest Sampling

To determine the effects of root exudation on soil metabolite composition and microbial recruitment during organic carrot production, both root exudates and soil were collected 10 weeks post-planting. At this point, carrots were in a vegetative, actively growing state, while exhibiting the phenotypic traits (e.g. color) of interest in this study, offering a suitable time to delineate potential differences in root exudation across genotypes. At sampling, 10 carrot roots were harvested from each plot. Rhizosphere soil was operationally defined as the soil attached to the carrot root after 5 seconds of gently shaking in the air (Wollum, A. G, 1994). Rhizosphere soil was gently removed to avoid damage to the roots and homogenized by plot. Bulk soil samples were collected by taking four soil cores per plot in between carrot rows to a 15 cm depth (5.7 cm diameter) and homogenizing by plot. Soil was transported to the lab on ice and roots were transported from the field to the lab in paper bags to maintain dark and warm conditions. Upon return to the lab, all soil samples were immediately sieved (2 mm) and a subsample was frozen at −80 °C for microbial community and metabolomics analysis.

To determine end-of-season impacts of carrot production on soil microbial community and metabolite composition, soil samples were taken concurrently with carrot harvest. At 14 weeks post-planting, carrots were harvested at maturity. Rhizosphere and bulk soil samples were collected and processed in an identical manner as the midseason samples.

### 2.4 Root Exudate Collection

Root exudates (n=20) were collected hydroponically after midseason harvest using methods adapted from Williams et al. (2021). First, three roots from each plot were randomly selected for exudate collection. Prior to root exudate collection, roots were gently rinsed of remaining soil using autoclaved deionized water and placed into acid washed and autoclaved glass beakers with 100 mL of molecular grade water as an extraction solution. Beakers were covered in aluminum foil to mimic dark growing conditions. Roots were submerged to allow for the diffusion of water-soluble metabolites into the water solution for 1.5 hours, which has been determined to be an appropriate amount of time for sufficient exudate collection while avoiding microbial turnover of exudate products (Neumann & Röhmheld, 2009). After roots were removed, the exudate solutions were immediately filter-sterilized through a 0.22µm filter, combined by plot, and frozen at −80°C. Tubes were lyophilized and the dried product was stored at −80°C prior to analysis.

### 2.5 Microbial Community Sample Preparation and Bioinformatics

DNA was extracted from 0.25g of rhizosphere and bulk soils collected at midseason and harvest sampling times using the DNeasy PowerSoil Kit (Qiagen, Germany) following manufacturer’s instructions, and the resulting purified DNA was stored at −80 °C. PCR reactions were performed using primer pairs that target the V4 region (~250 bp) of the 16S ribosomal RNA (rRNA) (hereafter “16S”) gene for bacterial communities (5’-3’; 515f GTGCCAGCMGCCGCGGTAA, 806r GGACTACHVGGGTWTCTAAT) as well as the internal transcribed spacer region 2 (hereafter “ITS2”) (~350-500 bp) for fungal communities (fITS7 GTGARTCATCGAATCTTTG, ITS4 TCCTCCGCTTATTGATATGC) (Ihrmark et al. 2012, White 1990, Turner et al. 1999, Kozich et al. 2013). 16S rRNA and ITS2 amplicon DNA was shipped on dry ice to Michigan State University’s Genomics Core (East Lansing, MI, USA) for library preparation and paired-end sequencing on a MiSeq (2 × 300 bp, Illumina, San Diego, CA, USA). Demultiplexed 16S and ITS2 sequences were processed using the DADA2 pipeline in R version 4.1.3 to construct amplicon sequence variants (Martin 2011, Callahan et al. 2016a, R Core Team 2022). Taxonomic classification was performed using the SILVA v138.1 and UNITE v9.0 databases for bacterial and fungal communities, respectively (McLaren & Callahan 2013, Abarenkov et al. 2023). Prior to analysis, sequences were rarefied to the minimum sequencing depth of 11925 and 9376 sequences per sample for 16S and ITS2 sequences, respectively in *phyloseq* (McMurdie & Holmes 2013, Callahan et al. 2016b). One 16S sample (from Block 3 H midseason rhizosphere) was removed from analysis due to poor sequencing results and one 16S sample (from Block 1 NR harvest rhizosphere) was identified as an outlier based on NMDS visual inspection and removed from analysis (see Supplementary Materials).

### 2.6 Root Exudate and Soil Metabolomics Data Acquisition

For untargeted metabolomics preparation and analysis, lyophilized root exudates and 10g of frozen soil samples were sent on dry ice to Colorado State University Proteomics and Metabolomics Facility (Fort Collins, CO, USA). Briefly, root exudates and soil samples were treated with a 50% or 80% MeOH-H_2_O solution, and then 1% formic acid was added to the soil samples. Samples were dried under a N_2_-stream, treated with 0.05 mL of methoxyamine hydrochloride in pyridine (25 mg/mL), and then treated with TMSTFA+1%TMCS for derivatization prior to analysis by gas chromatography-mass spectrometry (GC-MS; further detail in Supplementary Materials).

Prepared samples were injected into a Trace 1310 GC coupled to a Thermo ISQ mass spectrometer. Peak detection, alignment, and filling was performed using XCMS (version 4.2.2), and RAMClustR (version 1.2.4) was used to additionally normalize, filter, and group features (Smith et al. 2006, Broeckling, 2014). Feature matching was performed using the NIST 20 GC Method / Retention Index Database and the MS Dial GC-MS library (Babushok et al. 2007, Lai et al. 2017).

### 2.7 Univariate Data Analysis

Analysis of variance (ANOVA) was performed to test for significant effects and interactions of experimental factors Genotype (H, Hy, NR, PO), Time (Midseason, Harvest) and Soil Compartment (Bulk, Rhizosphere) on microbial richness based on the Chao1 Index (Chao 1984). When applicable, pairwise treatment means were compared with the Fisher’s Least Significant Difference (LSD; de Mendiburu 2023).

The package *ANCOMBC* (Analysis of Compositions of Microbiomes) was used to assess differentially abundant bacterial and fungal taxa across bulk and rhizosphere soils at the family level (Lin et al. 2020). Each genotype was analyzed at the midseason and harvest time point against its own bulk soil, using the bulk soil as a reference to assess differentially abundant taxa based on fold change values.

Metabolomics analysis was performed in Metaboanalyst 5.0 (Pang et al. 2021). To delineate which root exudates and soil metabolites were differentially abundant, one-factor ANOVA, fold change analysis and Wilcoxon t-tests were applied on each metabolite for every pairwise comparison of experimental factors using a fold change threshold of 1.5. Metabolites with a log2(fold change) value of >1.5 and a False Discovery Rate-adjusted p-value of <0.10 were considered significantly (<0.05) or marginally (<0.1) different.

### 2.8 Multivariate Analysis

To test for significant effects and interactions between carrot genotype, soil compartment, and time on microbial community composition, permutational analysis of variance (PERMANOVA) was performed on Bray-Curtis dissimilarities (Anderson 2001). A pairwise PERMANOVA post-hoc test was applied when appropriate. Similarly, PERMANOVA was used to distinguish the factors that significantly affected soil metabolome composition based on Euclidean distances. Multivariate homogeneity of group dispersions (beta dispersion) was similarly evaluated for microbial community compositions using the function ‘betadisper’ from *vegan* (Oksanen et al. 2022). PERMANOVA was carried out using the ‘adonis2’ function.

To assess associations between microbial community and metabolome dissimilarities, Mantel tests between distance matrices were performed in a pairwise manner using the ‘mantel’ function from *vegan* (Mantel, 1967; Oksanen et al. 2022). Bray-Curtis dissimilarity and Euclidean dissimilarity matrices were constructed for microbial community and metabolome datasets, respectively.

Distance-based redundancy analysis (dbRDA) was used to assess associations between metabolite concentrations and microbial community dissimilarity. To reduce data redundancy and the number of explanatory variables, root exudate and soil metabolites were subjected to a bottom-up clustering method wherein metabolites were clustered by similar behavior within the dataset using k-means clustering, such that metabolites with highly similar concentrations across all samples were reduced to a single variable. The minimum number of clusters was chosen such that within sums of squares was minimized, and that the means of each metabolite which were significantly different from the averaged value were minimized. Root exudate and soil metabolite clusters were then used as explanatory variables in dbRDA, and 16S and ITS communities were modeled separately. The significance of a cluster for explaining variation in microbial community dissimilarity in each dbRDA was assessed with a permutation test (999 permutations).

Raw sequence fastq files will be deposited into the NCBI SRA repository upon manuscript acceptance.

## 3. Results

### 3.1 Microbial Community Structure

Final feature tables contained 25350 (16S) and 5956 (ITS2) ASVs. The most abundant five phyla across all bacterial communities, representing nearly 70% of the total community, were Proteobacteria (26%), Actinobacteriota (14%), Acidobacteriota (13%), Firmicutes (9%) and Bacteroidota (7%). Likewise, the top three phyla across all fungal communities, representing 96% of the total community, were Ascomycota (59%), Mortierellomycota (19%), and Basidiomycota (18%). In total, 43 bacterial phyla and 16 fungal phyla were represented by at least one taxon.

Soil bacterial richness varied across genotypes and this effect was dependent on soil compartment (genotype × compartment, p < 0.05). Specifically, compared to their respective bulk soils, richness was increased by 19% in the rhizosphere of genotype Hy, and decreased by 10% in genotype PO (p < 0.05, Figure 1); neither genotypes NR nor H altered bacterial richness in their rhizosphere as compared to their respective bulk soils. Additionally, when comparing the rhizosphere across genotypes, bacterial richness was ~15% higher in genotypes H and Hy as compared to PO, and no other differences between genotypes were observed. When comparing the bulk soils across genotypes, bacterial richness also varied (p < 0.05). Specifically, bulk soil bacterial richness from plots planted with Hy was 14% lower than that of plots planted with NR. Further, the bulk soil of both Hy and NR were similar in richness to PO and H. Bacterial richness did not change across the two time points in either the rhizosphere or bulk soil.

**Figure 1.**
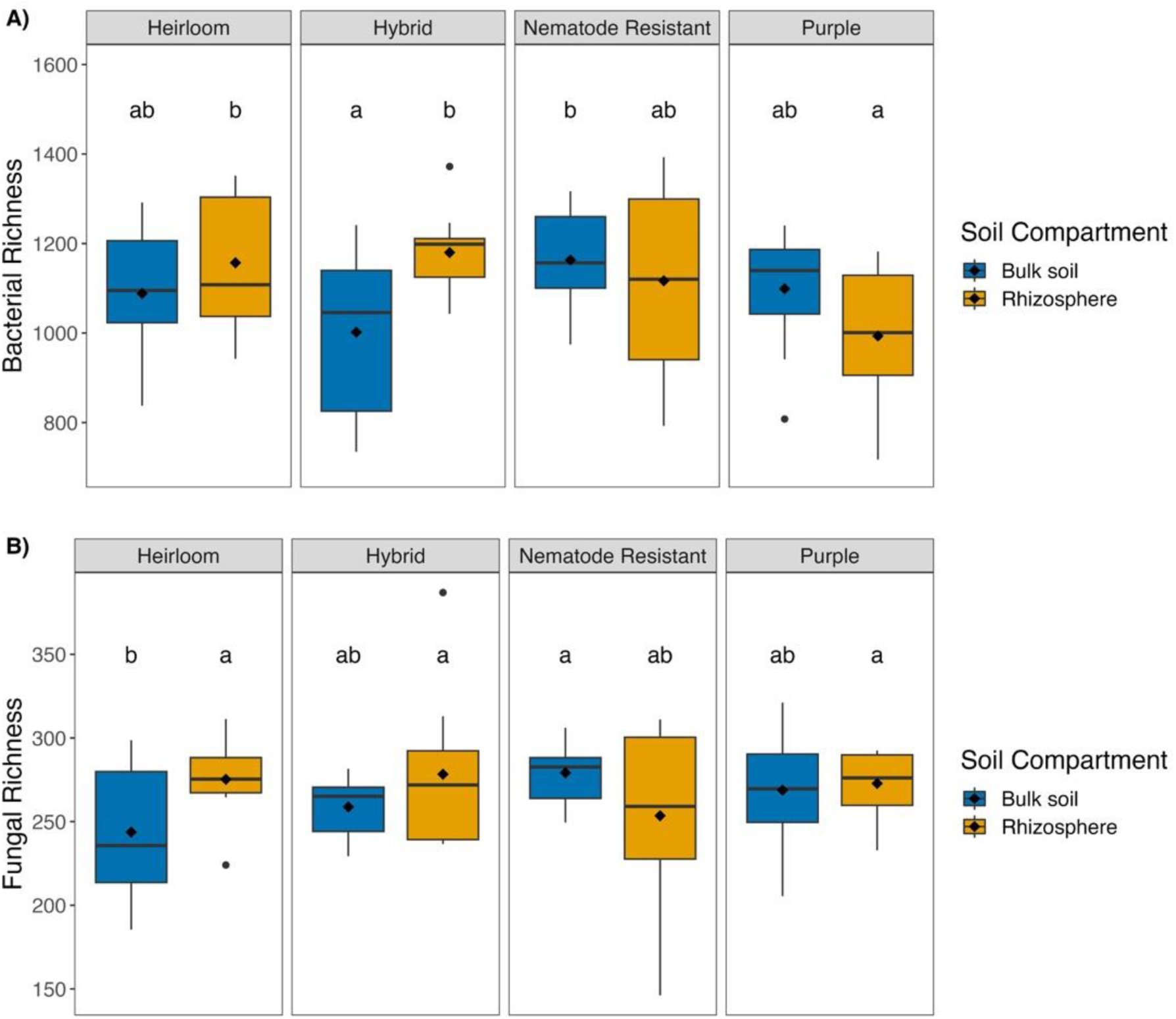
Boxplots demonstrate the differences in bacterial (A) and fungal (B) Chao1 richness between bulk and rhizosphere soils for each genotype. Medians are represented in the IQR region of the box as horizontal black lines, whereas diamonds represent the mean. Letters denote significant to marginally significant differences (p<0.1) between treatment means based on Fischer’s LSD.

Fungal richness was variably affected by soil compartment, genotype, and time (genotype × soil compartment, p <0.06, soil compartment × time, p < 0.05). Specifically, compared to its bulk soil, genotype H increased fungal richness by 13% in the rhizosphere, and no other genotypes significantly altered fungal diversity between bulk and rhizosphere soil (Figure 1). Across genotypes, fungal richness was similar in the rhizosphere, however, bulk soil richness varied. Specifically, bulk soil fungal richness from plots planted with genotype H was 9% lower compared to other bulk soils. Fungal richness was also 9% greater in the rhizosphere compared to the bulk soil at midseason, and this difference did not persist through harvest. From mid-season to harvest, fungal richness declined 8% in the rhizosphere, while bulk soil richness did not change overtime (Figure S1).

Bacterial community composition was distinct across genotypes, however, this effect was only present in the rhizosphere and not the bulk soil (PERMANOVA, genotype × soil compartment, p <0.10, Table 2, Figure 2). Across genotypes, bacterial communities also differed in the rhizosphere as compared to their respective bulk soils (pairwise PERMANOVA, p<0.05) and exhibited increased dispersion (betadisper, p<0.05). In the rhizosphere, when comparing across genotypes, bacterial communities were different across genotypes Hy, PO and NR, but not H, as such, H was similar to all genotypes. Across the mid-season and harvest time points, bacterial community composition was distinct in both rhizosphere and bulk soils (pairwise PERMANOVA, p<0.05).

**Figure 2.**
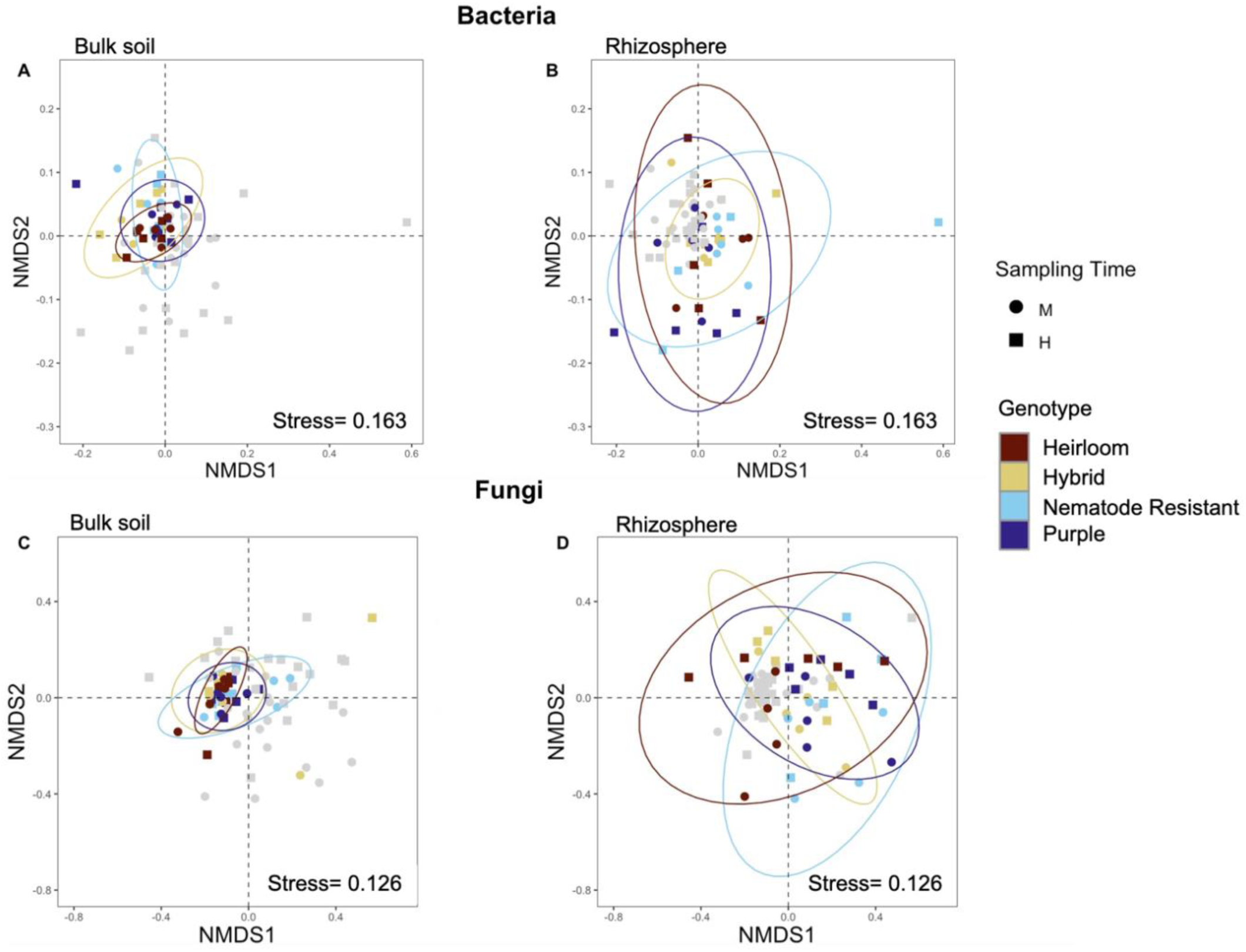
Global visualization of bacterial (A, B) and fungal (C, D) β-diversity and dispersion. Non-metric multidimensional scaling was performed on Bray-Curtis dissimilarities at three dimensions and visualized using axes 1 and 2. Left: Bulk soils in the ordination are highlighted, and ellipses show the 95% confidence interval for each genotype. Right: Rhizosphere soils in the ordination are highlighted, and ellipses show the 95% confidence interval for each genotype. M, Midseason, H, Harvest.

**Table 2.** Results of PERMANOVA tests of experimental factors on bacterial and fungal community composition. P-values for which post-hoc tests were performed on the associated factor or interaction are bolded. G, Genotype; SC, Soil Compartment; T, Time.

|  | Bacteria |  | Fungi |  |
| --- | --- | --- | --- | --- |
|  | R<sup>2</sup> | P | R<sup>2</sup> | P |
| Genotype | 0.040 | 0.043 | 0.043 | **0.047** |
| Soil Compartment | 0.029 | 0.002 | 0.056 | 0.001 |
| Time | 0.016 | 0.001 | 0.033 | 0.001 |
| Block | 0.025 | 0.001 | 0.027 | 0.002 |
| G:SC | 0.040 | **0.061** | 0.04 | 0.113 |
| G:T | 0.036 | 0.715 | 0.033 | 0.481 |
| SC:T | 0.016 | **0.002** | 0.023 | **0.005** |
| G:SC:T | 0.038 | 0.303 | 0.038 | 0.184 |

Fungal communities differed in the rhizosphere as compared to the bulk soil (PERMANOVA, soil compartment, p<0.01, Table 2, Figure 2), and exhibited increased dispersion (betadisper, p<0.05). When comparing across genotypes, independent of soil compartment, fungal communities in plots planted with genotype NR were different from plots planted with H (PERMANOVA, p<0.05, Table 2). Similar to that observed with the bacteria, fungal community composition was unique in both rhizosphere and bulk soils across the mid-season and harvest time points (PERMANOVA, soil compartment × time, p< 0.01, Figure 2).

### 3.2 Rhizosphere Microbial Recruitment

To determine whether root exudates selectively recruit microbial taxa in the rhizosphere, we categorized differentially abundant microbial families as positive or negative responders to root growth in a framework adapted from Zhalnina et al. (2018). Bacterial recruitment at the family level was distinct across genotypes and across time, though this trend was driven by differences in abundance of relatively few taxa (Figure 3). Specifically, *Rhodocyclaceae* was the only taxon to display temporal consistency, and positively responded to genotype NR at both midseason and harvest. Additionally, *Nitrsosphaeraceae* responded positively to both genotype NR and H at midseason. At midseason, the top positive responder for each genotype was order *Kapabacteriales* (family unclassified), *Rhodocyclaceae,* order SJA-28 (family unclassified), and *Methylophilaceae* for genotypes Hy, NR, PO and H, respectively. At harvest, the top positive responder for each genotype was *Methylophilaceae*, *Flavobacteriaceae*, *Micrococcaceae*, and *Sphingobacteriales*, for genotypes Hy, NR, PO and H, respectively. Conversely, the top negative responders for each genotype at midseason were *Chthonomonadaceae*, phylum Chlorofexi (family unclassified), *Anaeromyxobacteraceae,* and phylum NB1-j (family unclassified), for genotypes Hy, NR, PO and H, respectively. At harvest, no negative responders were identified for genotype Hy. Families *Peptostreptococcaceae*, order *Polyangiales* (family unclassified) and order *Bacillales* (family unclassified) were top negative responders for genotypes NR, PO and H, respectively.

**Figure 3.**
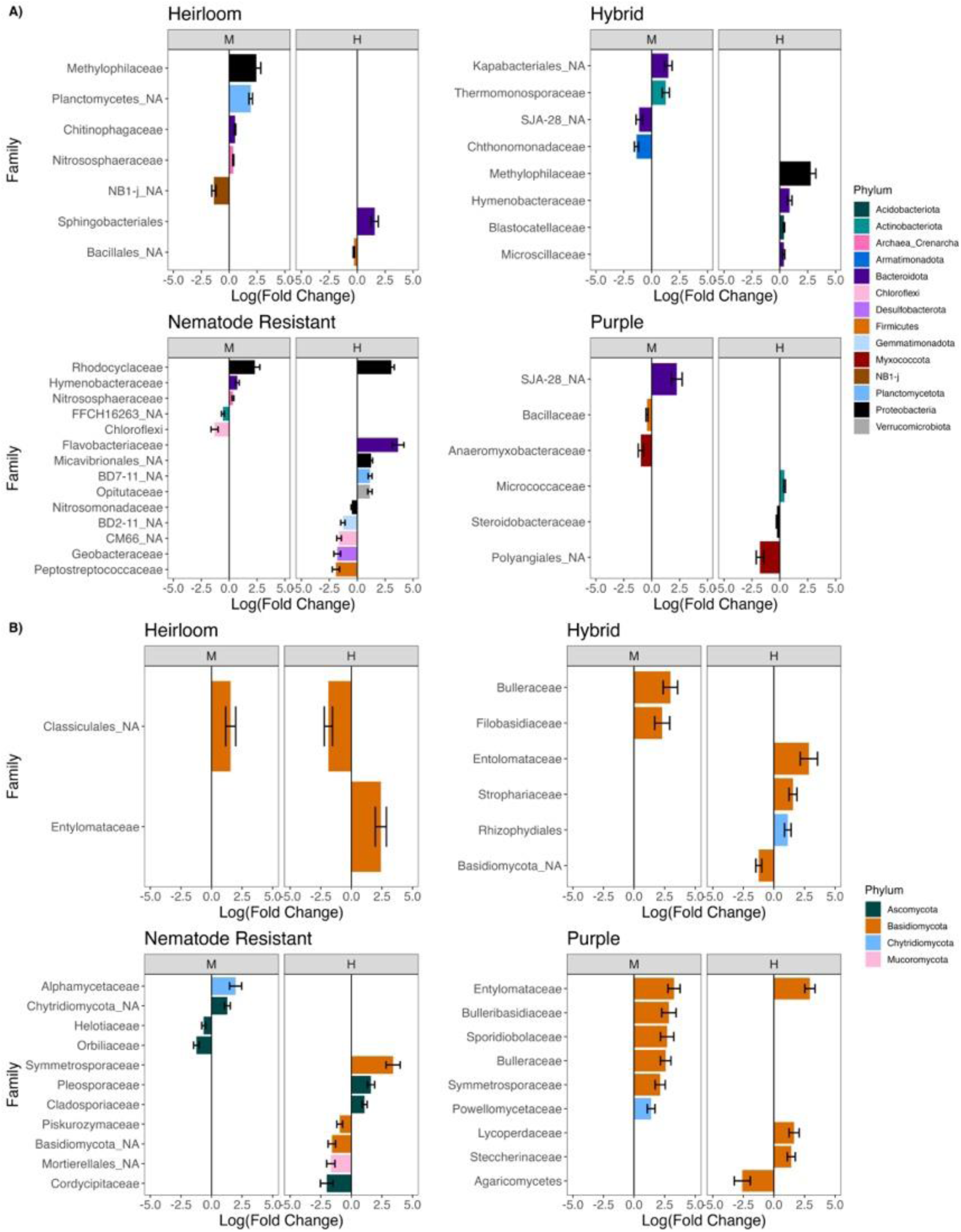
For each carrot genotype, positive and negative responding bacterial (A) and fungal (B) recruitment in the rhizosphere are mapped across midseason (M) and harvest (H) time-points. Negative log(fold change) values represent taxa of greater relative abundance in the bulk soil, and positive log(fold change) values represent taxa of greater relative abundance in the rhizosphere of the respective genotype. Taxa with NA in their name represent unidentified family-level classification, and the next available classification level is given.

Fungal recruitment was also distinct across genotypes and time (Figure 3). At midseason, the top positive responder for each genotype was *Bulleraceae*, *Alphamycetaceae, Entolomataceae*, and class *Classiculales* (family unclassified) for genotypes Hy, NR, PO and H, respectively. At harvest, the top positive responder for each genotype was *Entolomataceae* for genotypes Hy and H, *Symmetrosporaceae* for genotype NR, and class *Agaricomycetes* for genotype PO. *Orbiliaceae* negatively responded to genotype NR at midseason. *Basidiomycota, Symmetrosporaceae Agaricomycetes* and *Entolomataceae* were top negative responders at harvest for genotypes Hy, NR, PO and H, respectively.

### 3.3 Root Exudate and Soil Metabolite Composition

To determine whether carrot genotypes exuded different compositions of low-molecular weight compounds and if soil metabolites were differentially abundant between genotypes, 128 and 114 unique compounds were annotated for the root exudate and soil metabolite extractions, respectively. Of those, 33 root exudates and 49 soil metabolites were identified by our database search. Notably, three unidentified root exudates (C056, C063, C115) made up roughly 50% of the total spectral abundance of exudates across all genotypes, and 20-30% of the total spectral abundance was composed of unidentified, low abundance compounds (see Supplemental Materials).

No root exudates were unique between genotypes, however, 37 compounds were differentially abundant across genotypes. These included sucrose, citric acid, psicose, malic acid, ketohexose, galactose, quinic acid, scyllo-inositol, methyl 2,6-dihydroxybenzoate (for simplicity, hereafter “benzoate”), alpha-(4-dimethylaminophenyl)-omega-(9-phenanthryl)octane (hereafter “octane”), silane-diethylisohexyloxy(3-methylbutoxy; hereafter “silane”), as well as 30 unidentified compounds. Of these 37 compounds, 34 were most abundant in genotype H (Figure 4). Conversely, genotype NR consistently had a lower abundance of most compounds compared to other genotypes. Of the identified compounds, genotype Hy was more abundant in benzoate and scyllo-inositol. Genotype NR had a lower abundance of all compounds except for citric acid, which was lowest in genotypes H and PO. Genotype NR was relatively low in sucrose as compared to other genotypes; sucrose was relatively most concentrated in genotypes PO and H (Figure 4).

**Figure 4.**
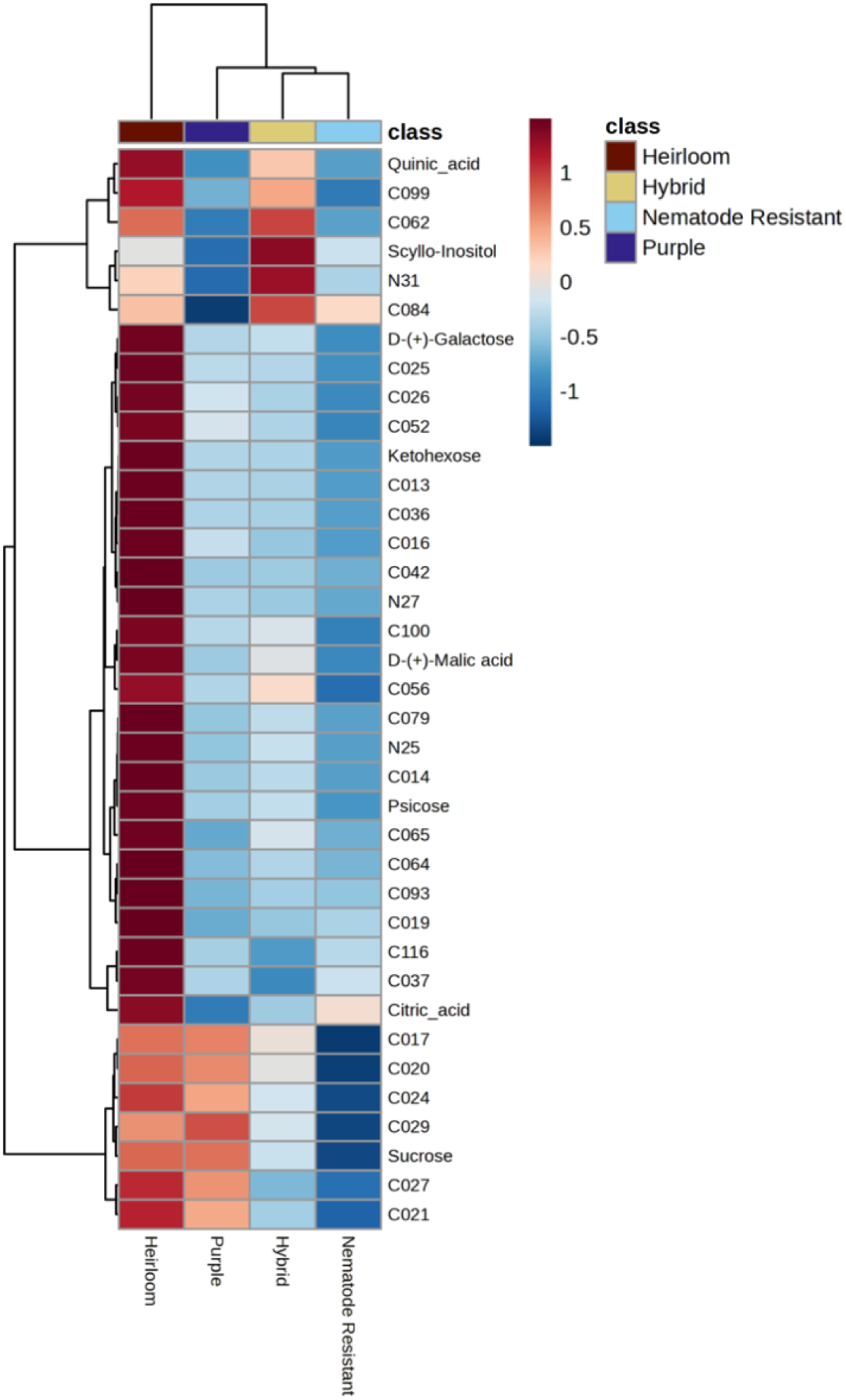
Differential abundance of root exudates across genotypes. The spectral intensity of compounds was scaled based on deviation from the mean using z-scores, clustered, and grouped by genotype to illustrate up or down expression. N25, “octane”; N27, “silane”; N31, “benzoate”. Figure was constructed in Metaboanalyst 5.0.

Soil metabolite composition was different in the rhizosphere compared to the bulk soil, and was different across time, but did not differ between genotypes (PERMANOVA, soil compartment, p<0.05, time, p<0.05). Two unidentified compounds were highly different between soil compartments; specifically, compound C041 was highly abundant in the rhizosphere and C076 was highly abundant in the bulk soil. More soil metabolites shifted across time than in response to soil compartment, and several compounds were differentially abundant at the midseason or harvest sampling point (Figure S2, Table S3).

### 3.4 Microbe-Metabolite Relationships

Root exudates and soil metabolites were differentially associated with fungal and bacterial communities (Table 3). Specifically, bacterial, but not fungal community composition was significantly associated with root exudate chemical composition (mantel test, p<0.05) and was marginally associated with soil metabolite composition (p<0.1). Moreover, fungal community dissimilarity was not significantly associated with the compositional dissimilarity of either root exudates or soil metabolites (Table 3).

**Table 3.** Results of Mantel tests between soil microbial and metabolite dissimilarities.

|  | Bacteria |  |  |  | Fungi |  |  |  |
| --- | --- | --- | --- | --- | --- | --- | --- | --- |
|  | Pearson |  | Spearman |  | Pearson |  | Spearman |  |
|  | r | P | r | P | r | P | r | P |
| Soil Metabolome | 0.1112 | 0.109 | 0.1193 | 0.086 | 0.019 | 0.391 | 0.037 | 0.318 |
| Root Exudates | 0.2409 | 0.043 | 0.236 | 0.037 | 0.010 | 0.47 | -0.002 | 0.504 |

To assess if and how root exudate and soil metabolite concentrations explained soil microbial community composition, root exudate and soil metabolites were each first clustered into six groups based on similarity in spectral intensity using k-means clustering. Root exudate concentrations accounted for a significant proportion of the variation in rhizosphere bacterial (35%), but not fungal community dissimilarity (Figure 5). Conversely, clustered soil metabolites accounted for a significant portion of fungal (11%), but not bacterial community dissimilarity (Figure 6). Root exudate concentrations were not significantly associated with soil metabolite composition in the rhizosphere. Root exudate clusters, specifically cluster 2 (compounds C056, C063, C115) and cluster 6 (oxalic acid), were significantly associated with rhizosphere bacterial communities across genotypes (Figure 5). In the constrained ordination, cluster 2 presented a positive association across axis 1 and 2, and was positively associated with rhizosphere microbial community dissimilarity of genotype Hy. Oxalic acid was positively associated with axis 1 and negatively associated with both axis 2 and rhizosphere microbial community dissimilarity of genotype NR (Figure 5). Among soil metabolites clusters, cluster 4 and cluster 5 were significantly associated with fungal community composition. In the constrained ordination, soil cluster 4 was positively associated with axis 1 and 2 as well as with midseason fungal communities, and more so of those in the rhizosphere. Soil cluster 5 had a strong positive association with axis 2 and was positively associated with rhizosphere fungal communities at harvest (Figure 6).

**Figure 5.**
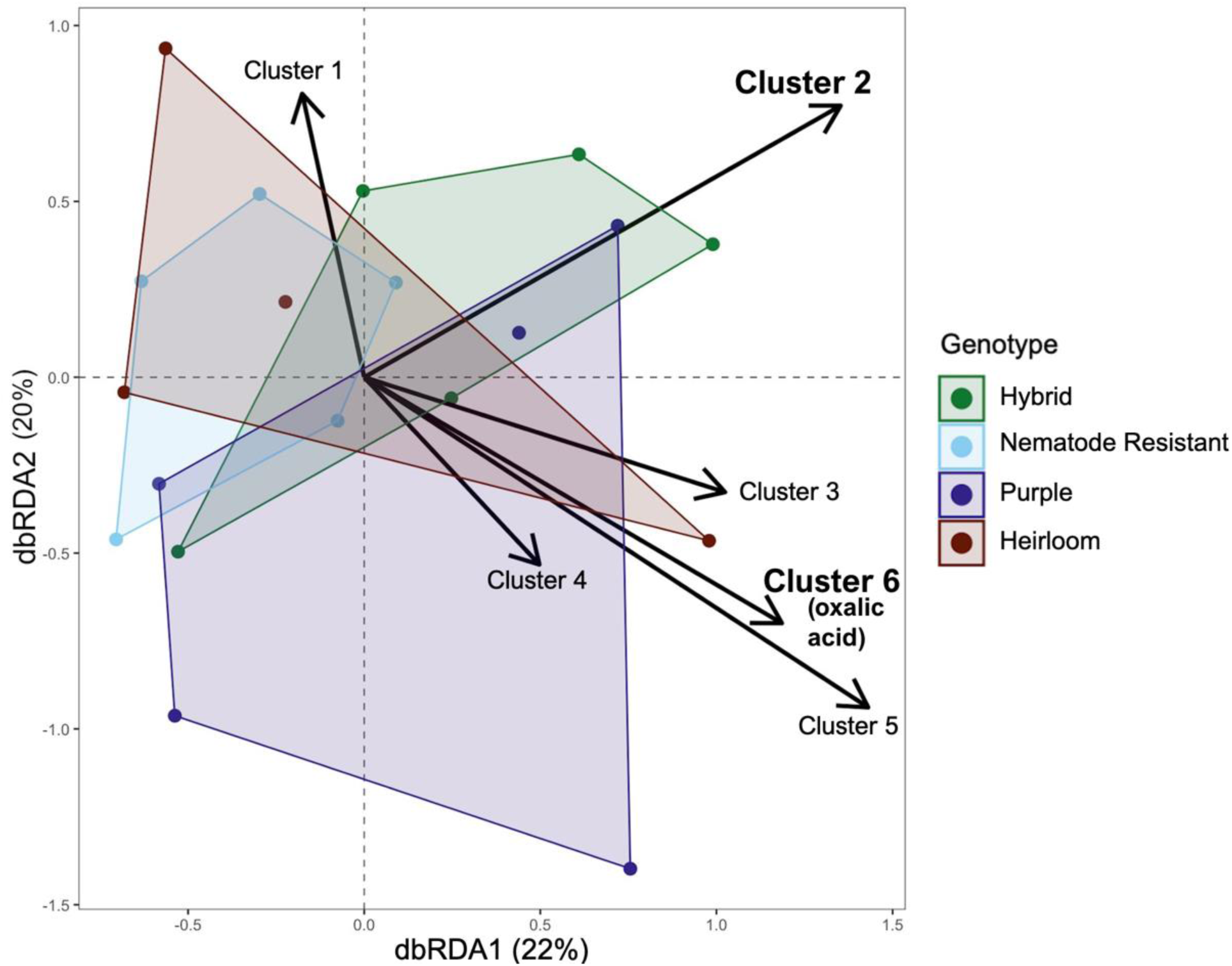
Distance-based redundancy analysis of root exudate clusters on midseason rhizosphere bacterial communities. Root exudate clusters were modeled against a Bray-Curtis dissimilarity matrix of bacterial communities. Clusters 2 and 6 (large text) significantly explained fungal community composition. Bacterial communities are colored by genotype, and polygons represent the convex hulls of each geometric distribution of points on the ordination within a genotype.

**Figure 6.**
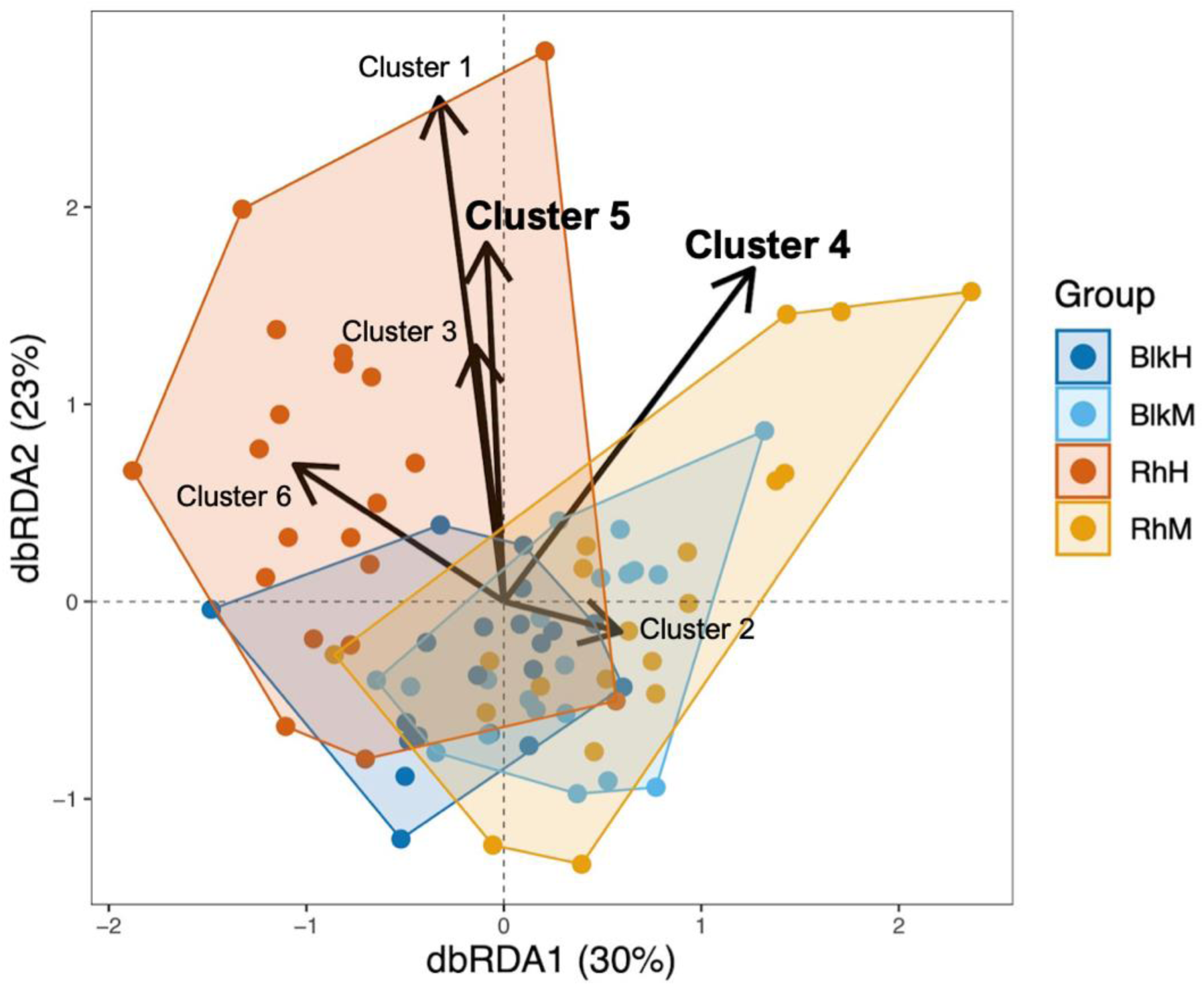
Distance-based redundancy analysis of soil metabolite clusters on soil fungal communities. Soil metabolite clusters were modeled against a Bray-Curtis dissimilarity matrix of fungal communities. Clusters 4 and 5 (large text) significantly explained fungal community composition. Fungal communities are colored by soil compartment and time, and polygons represent the convex hulls of each geometric distribution on the ordination of points within a group. BlkH, Bulk soil at harvest; BlkM, Bulk soil at midseason; RhH, Rhizosphere at harvest; RhM, Rhizosphere at midseason.

## 4. Discussion

Across four phenotypically diverse genotypes of carrot, we determined the influence of novel trait breeding on root exudate, soil metabolite and soil microbial community composition. Including two established and two novel genotypes, we tested the hypothesis that root exudation mediates microbial rhizosphere recruitment, and that genotype-specific shifts in root exudate composition would alter microbial communities and soil metabolite composition. In support of hypothesis 1.1, genotypes differentially exuded low-molecular weight compounds and hosted compositionally different microbial communities (Figures 2, 4). In support of hypothesis 1.2, microbial recruitment from a bulk soil to the rhizosphere soil was distinct across genotypes and across the growing season (Figure 3). Hypothesis 2.1 was not supported, as root exudate and soil metabolite composition were distinct and were not associated with one another (Table 3). Finally, hypothesis 2.2 was partially supported, as root exudates were significantly associated with bacterial and not fungal communities, while soil metabolites were only weakly associated with fungal and bacterial communities (Figures 5, 6, Table 3).

### 4.1 Root exudation & soil metabolites exert a differential influence on bacterial and fungal communities

Root exudates were significantly associated with bacterial, but not fungal communities, suggesting that bacteria are the primary responders to compositional differences in low molecular weight compounds from root exudates in the rhizosphere. Additionally, root exudate chemical composition explained 35% of the variation across the rhizosphere bacterial communities while categorical differences in genotypes only explained 4% of the variation (Figure 5). These observations differ from previous work on carrots in a greenhouse setting with artificial soil, where the carrot genotype accounted for 50% of bacterial variation (Triviño 2023), alluding to the need to better understand the mechanisms shaping rhizosphere bacterial communities in field soil.

Our results are indicative of a shared root exudate metabolome across carrot genotypes where differences in the concentration of individual compounds, rather than the presence of unique compounds among genotypes, ultimately shape the composition of the rhizosphere bacterial community. Indeed, the compounds with the strongest associations with bacterial community compositional dissimilarity were highly abundant across all genotypes. Notably, oxalic acid was significantly associated with bacterial community composition and is universally recognized for its role in plant nutrient acquisition through the solubilization of phosphorus (Bolan et al. 1994). Oxalic acid was previously reported as a major organic acid in root exudates of horticultural crops (Xiang et al. 2020, Ling et al. 2011, Vančura & Hovadik 1965). Additionally, the correlation between oxalic acid concentration and bacterial community composition aligns with previous observations that organic acids can be highly explanatory of bacterial community shifts. (Ulbrich et al. 2022, Landi et al 2006, Shi et al 2011). This indicates that genotype-specific differences in oxalic acid production, and other currently unidentified compounds, may affect bacterial community composition in the carrot rhizosphere.

Soil fungal communities were more strongly associated with the soil metabolome as compared to the root exudate metabolome. Fungal communities in the rhizosphere were distinct from the bulk soil, though there were no effects across genotypes, and fungal communities were not well-explained by root exudate composition. Based on these results, fungal recruitment may be more mediated by differences in soil carbon availability than specific metabolic capabilities, as compared to bacteria. Though, soil fungal communities did differ from mid-season to harvest, and these changes were associated with variation in soil metabolite concentration (Figure 6), suggesting that fungal metabolism is more influenced by seasonal conditions than plant genotype. Some of the low molecular weight compounds identified in the soil compartment have been reported to be produced by fungi, suggesting that they are of microbial and not plant-derived origin. Specifically, the aromatic compound danthron can be produced by some members of the Ascomycota (Anisha et al. 2018). Further, an isolate of *Fusarium* has been reported to produce rhein (You et al. 2013). Both of these compounds are derivatives of anthraquinone, which is produced by fungi across systems and possesses pathogen suppressive qualities (Gessler et al. 2013, Fouillaud et al. 2016, Masi et al. 2020). These compounds were members of soil metabolite clusters four and five, which were significantly associated with fungal community composition (Figure 6). This suggests certain fungi can modify the soil metabolome in agricultural settings through the production of danthron, rhein, and possibly several unidentified compounds. Additionally, greater soil concentrations of glucose, maltotriose and trehalose at harvest suggest a shift in fungal carbohydrate metabolism throughout the growing season (Table S3). To further explore this possibility, potential links between bulk soil composition, fungal communities, and the soil metabolome could be more closely interrogated.

Interestingly, metabolite profiles from rhizosphere soil were neither significantly correlated with root exudate metabolite composition nor bacterial or fungal community composition. These observations are counter to the prediction that root exudate composition would directly modulate metabolite presence in the rhizosphere and govern rhizosphere microbial composition (Bi et al. 2020, Song et al. 2020). It is possible this is because root exudates are directly and quickly assimilated by rhizosphere bacteria and are not directly incorporated into the rhizosphere metabolome. It has been previously observed that root exudates only diffuse into a small area, with estimates of diffusion from 2 to 10 mm of soil away from the root (Sauer et al. 2006, Raynaud 2010). This result is consistent with the hypothesis that root exudates are rapidly utilized by microorganisms and have little influence on soil metabolomes. Unique profiles of soil metabolites as compared to the plant-derived metabolites present in the root exudates in this study indicates that soil metabolome composition is more greatly shaped by bulk soil composition and microbial production rather than direct plant inputs. Notably, there is a lack of literature evaluating root exudates and soil metabolites in tandem as we present here, and more research is needed to understand if there is a relationship between the two at various spatial scales in the rhizosphere.

### 4.2 Root exudation is distinct across genotypes and most different between heirloom and nematode resistant genotypes

While root exudate profiles were similar in membership overall across genotypes, abundances of certain compounds were strikingly different among genotypes. Most notably, genotype H, an heirloom genotype, produced greater abundances of several carbohydrates, organic acids and numerous unidentified compounds compared to other genotypes, whereas genotype NR produced less of these compounds as compared to other genotypes (Figure 4). While underexplored in vegetables, domestication of other crops shifts root exudate profiles, suggesting that differences between genotypes are a consequence of breeding history, and that older genotypes may maintain distinct root exudation patterns as compared to more heavily bred genotypes (Yue et al. 2023). The heirloom genotype H has been previously reported to respond more positively to root colonization of arbuscular mycorrhizal fungi, compared to hybrid genotypes (Pearson et al. 2020). Our results provide further evidence that breeding history could shift plant-microbe interactions in a vegetable system.

In this study, the heirloom genotype H had higher exudation of organic acids and carbohydrates, perhaps indicative of greater resource investment in microbial stimulation than other genotypes. This result is consistent with the emerging hypothesis that heirloom genotypes benefit more from plant-soil interactions than highly bred genotypes that prioritize yield (Pearson et al. 2020). Carbohydrates are notably consequential for soil microbes, especially bacteria, in the rhizosphere due to their abundance in root exudates. Specifically, they have been associated with enhanced microbial activity, and have been linked to greater phytohormone production in some plant-associated bacteria as well as greater mineralization of soil organic matter when root exudate-nitrogen is low, harboring potential benefits for plant growth (Lloyd et al. 2016, Seitz et al. 2022).

Additionally, in this study, a distinct root exudate profile was observed from the nematode resistant genotype NR relative to other genotypes (Figure 4). While nematode resistance was not assessed in this study, relationships between these two traits should be further explored, especially as nematode presence can influence microbial carbon and nitrogen cycling through trophic interactions (Kane et al. 2023). Our results suggest that novel trait introduction, in addition to breeding history of genotypes, might shift root exudate composition, and more specific links between root exudates and traits should be investigated.

### 4.3 Bacterial recruitment in the rhizosphere is distinct across genotypes and suggests differential consequences for soil biological functioning

Bacterial selection in the rhizosphere from a bulk soil varied across genotype, root exudate composition, and time-point, providing support that microbial recruitment is associated with plant developmental stage and phenotypic identity (Chaparro et al. 2018, de Ridder-Duine et al. 2005, Micallef et al. 2009, Aira et al. 2010). This genotype-mediated diversification of soil bacterial communities through root exudation might indicate consequences for community functional capacities (Figure 3). For example, *Rhodocyclaceae* was selectively enriched in genotype NR at both midseason and harvest, and has previously been identified as a denitrifying bacteria, as it contains *nirS* (Yu et al. 2018). *Nitrsosphaeraceae* was positively associated with genotypes NR and H; *Nitrsosphaeraceae* is an ammonia oxidizing bacteria, which are considered indicators of soil health (Mundepi et al. 2019). These results align with observations that genotype NR hosts distinct N-associated functions (e.g., ammonia oxidation; Triviño et al. 2023). Genotype H was additionally enriched with members of the *Chitinophagaceae*, a family identified as containing N fixing taxa, at midseason (Martin et al. 2022). *Methylophilaceae*, a methylotroph, was another notable taxon that was enriched in genotype Hy at midseason and genotype H at harvest. Methylotrophs have been previously identified to utilize root exudates in the rhizosphere of pea and wheat (Macey, 2017). Further, they have been reported to synthesize phytohormones including auxin and cytokinins, and their role as plant-growth promoters is being explored for agricultural uses (Ponnusamy et al. 2017, Ivanova et al. 2001).

Genotype NR had lower abundances of most root exudates (Figure 4), and previous research has identified this genotype as having relatively higher nitrogen use efficiency, possibly through the selection of specific bacterial taxa. In contrast, genotype H had higher abundances of most root exudates, suggesting a larger investment in microbial stimulation overall. These results suggest that across genotypes, there is a general trend towards functional redundancy and selection for N-related functions, however, the mechanisms of this selection across genotypes may be different.

### 4.4 Limitations and Future Directions

While we used a GC-MS based approach for this study that is coarser in resolution than other technologies used in metabolomics inquiries, differences between genotypes may be discriminated more strongly by employing methods such as high-resolution accurate mass LC-MS that could further characterize compounds that express genotype-specific patterns. Additionally, while our results coincide with previous findings that heirloom and nematode resistant genotypes express distinct characteristics, the inclusion of multiple genotypes of varying ancestral lineage or degrees of nematode resistance could strengthen this line of inquiry. To bridge links between plant genetics and root exudation, transcriptomic evaluation of carrot gene expression to characterize the functional genomics underlying exudate composition should also be explored. Finally, more quantitative measures of root exudation rates and soil microbial processes could provide valuable insights into genotype-mediated shifts in soil biological outcomes.

### 4.5 Conclusions

The results of this study suggest that plant-soil microbe interactions are influenced by trait breeding in organic carrot. This aligns with previous findings that carrot genotypes host distinct root-associated microbiomes that potentially differ in their functionality (Triviño et al. 2023, Abdelrazek et al. 2020). Moreover, we show that root exudation patterns were distinct across genotypes. We provide further evidence that selection for microbial taxa with distinct N-related functions is influenced by genotype identity, and that this selection could explain previously demonstrated differences in nutrient use efficiency in novel genotypes. Further, the heirloom genotype potentially enriched plant-growth promoting bacterial activity in the rhizosphere through exuding relatively greater levels of carbohydrate and organic acid production and selecting for taxa with demonstrated potential for phytohormone production and nitrogen fixation, relative to genotypes of contrasting breeding histories. These results indicate that both breeding history and novel trait introduction in plant genotypes can diversify plant-microbe interactions and resulting soil biological functional capacities.

Root exudation was significantly associated with bacterial communities, suggesting that breeding for specific root exudate profiles to target and manipulate rhizosphere bacteria could be a strategy for plant or soil health objectives. However, more research is needed to understand the drivers of plant-fungal relationships and potential applications for organic agriculture, especially as strengthening beneficial fungal associations is a notable priority in organic agroecosystems. Through the investigation of plant-microbe interactions, we show that cultivar development that addresses grower needs can occur in tandem with considerations for soil biological functioning, supporting a systems-based philosophy in agricultural practices.

## Funding and Acknowledgements

This material is based upon work that is supported by the National Institute of Food and Agriculture, U.S. Department of Agriculture Grant No. 5090-21000-073-066-A. We gratefully acknowledge the research contributions of the West Madison Agricultural Research Station Staff, members of the Silva and Dawson labs, Dr. Michelle Pearson, Annalise Keaton, Gabriela Martinez and Kathryn Lindeman. Thank you to the Colorado State University Analytical Resources Core and the Michigan State University Genomics Research Technology Support Facility with assistance with metabolomic and microbiome analysis.

## Declaration of competing interest

The authors declare no conflict of interest.

## Supplementary Materials

**Figure S1.**
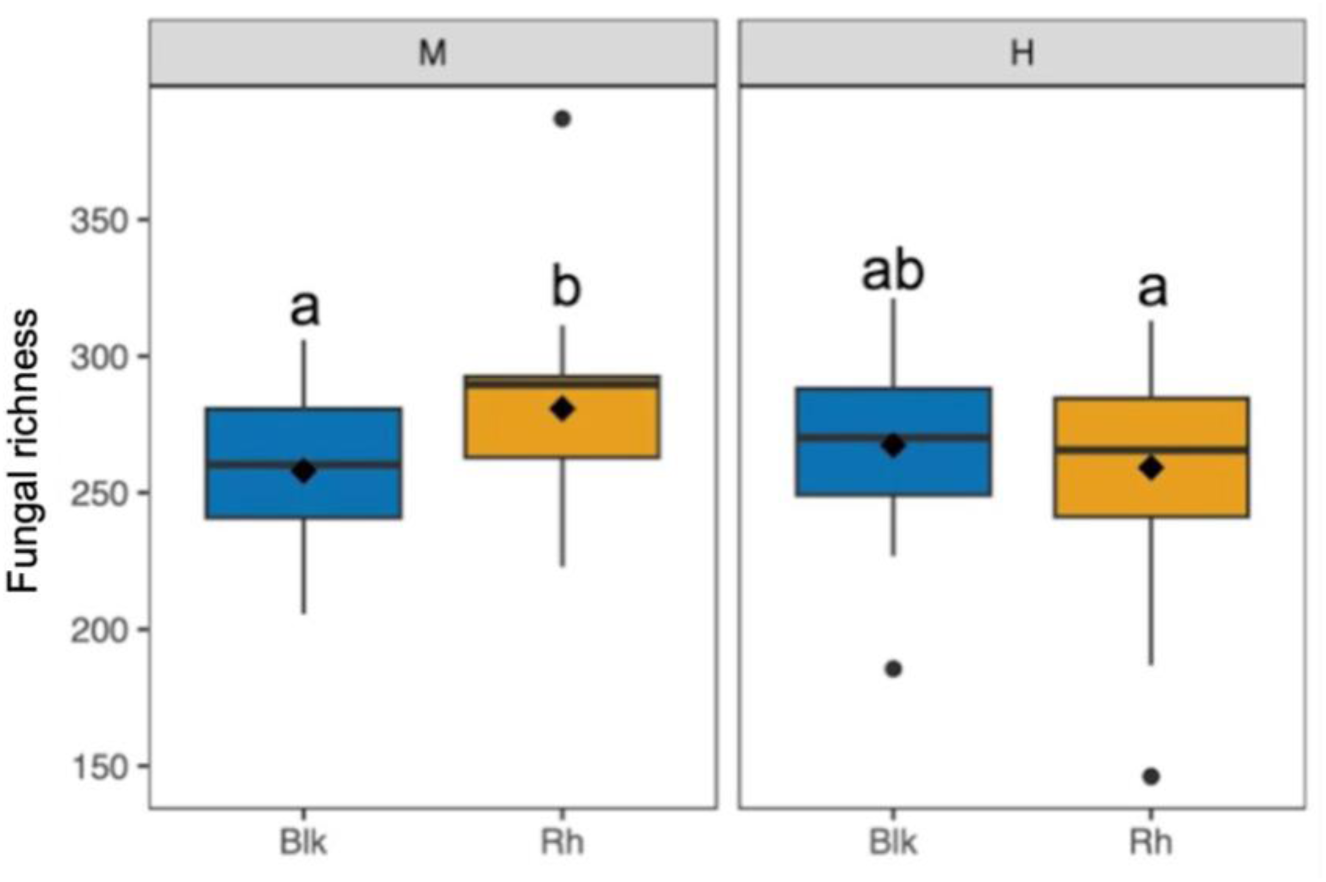
Fungal richness at each sampling timepoint. Boxplots demonstrate the differences in Chao1 richness between bulk and rhizosphere soils. Medians are represented in the IQR region of the box as horizontal black lines, whereas diamonds represent the mean. Letters denote significant to marginally significant differences (p<0.1) between treatment means based on Fischer’s LSD. M, Midseason; H, Harvest, Blk, Bulk soil, Rh, Rhizosphere.

**Figure S2.**
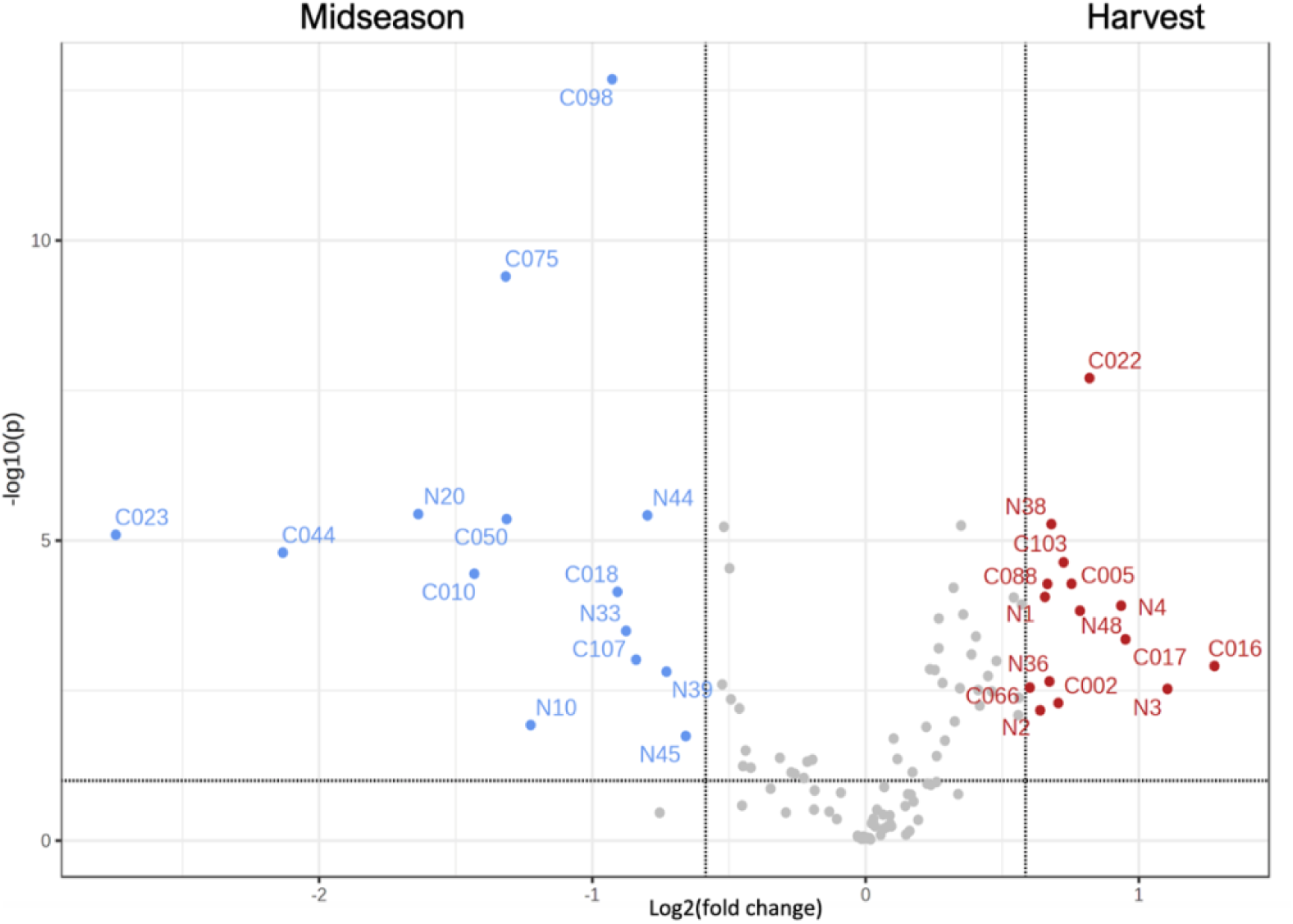
Differential abundance of soil metabolites across sampling timepoints. See Table S3 for metabolite identities. Plot was constructed in Metaboanalyst 5.

**Figure S3.**
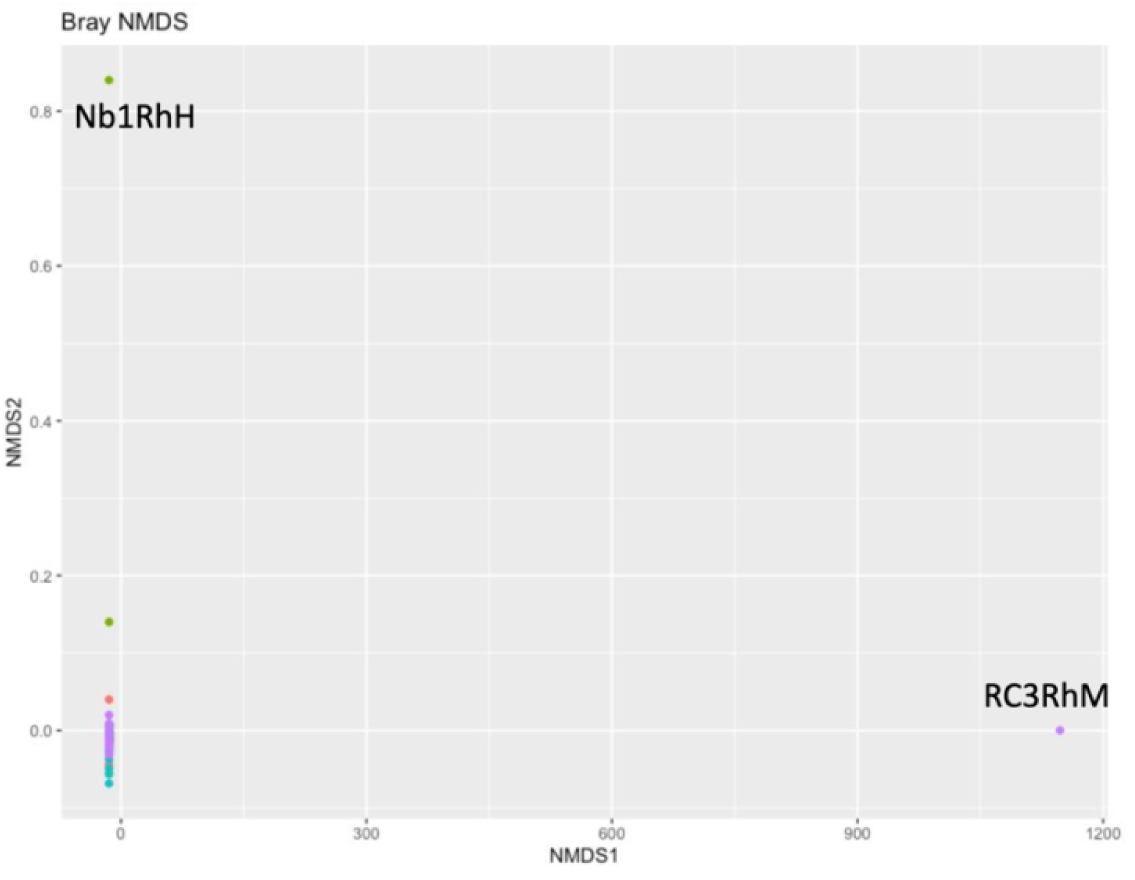
Visual inspection of NMDS plot with outliers. (RC3RhM; Red Core Chantenay Block 3, Rhizosphere at Midseason was removed due to extremely poor sequencing results. Nb1RhH; Nb8503 Block 1, Rhizosphere at Harvest was removed due to unusual results as visualized in the NMDS plot).

**Table S1.**
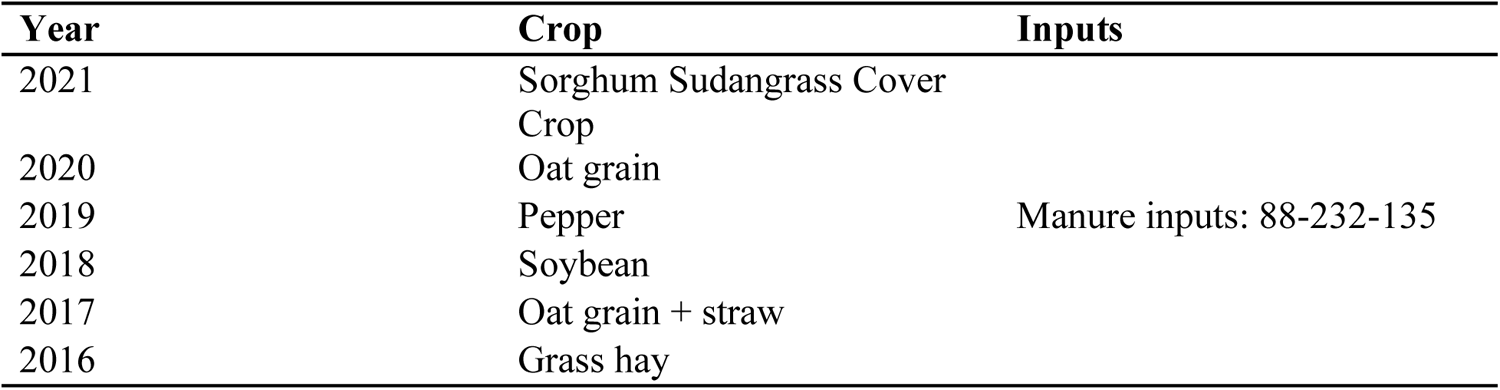
Crop rotation history for experimental plots. Acquired from West Madison Agricultural Research Station Staff.

**Table S2.** Soil test history for experimental plots. Acquired from West Madison Agricultural Research Station Staff.

| Soil Name | Kegonsa series |  |  |  |  |
| --- | --- | --- | --- | --- | --- |
| Soil Texture | Silt loam |  |  |  |  |
| Slope | 4% |  |  |  |  |
| Test Date | Plow Depth (in) | Avg pH | Avg OM (%) | Avg P (ppm) | Avg K (ppm) |
| 10/22/2021 | 6 | 7.3 | 3 | 52 | 185 |
| 10/19/2020 | 6 | 7.1 | 3.4 | 59 | 145 |
| 12/11/2018 | 7 | 7.1 | 2.8 | 54 | 143 |
| 11/15/2017 | 7 | 7.1 | 2.3 | 29 | 94 |
| 11/22/2016 | 7 | 6.8 | 2.9 | 28 | 123 |
| 12/2/2015 | 7 | 7.4 | 2.5 | 59 | 220 |
| 12/6/2013 | 7 | 6.9 | 2.7 | 43 | 179 |

**Table S3.** Identified soil metabolites referenced in Figure S2.

| Compound name | Code | Up-expression |
| --- | --- | --- |
| Cyclohexasiloxane, dodecamethyl | N10 | Midseason |
| 2,3-Dimethylquinizarin, bis(trimethylsilyl) ether | N20 | Midseason |
| 4-Hydroxyanthraquinone-2-carboxylic acid | N33 | Midseason |
| 3-Hydroxyflavone, triethylsilyl ether | N39 | Midseason |
| Silane, diethylhexadecyloxy(2-methoxyethoxy) | N44 | Midseason |
| Rhein | N45 | Midseason |
| D-Trehalose | N1 | Harvest |
| DL-beta-Hydroxybutyric acid | N2 | Harvest |
| Glucose_1 | N3 | Harvest |
| Maltotriose | N4 | Harvest |
| 1-Isopropoxy-2-trihexylsilyloxybenzene | N36 | Harvest |
| Norleucine | N38 | Harvest |
| Methyl 2-Trimethylsiloxy-octadecanoate | N48 | Harvest |

### Additional Metabolomics Methodology

#### Root exudates

Each sample in the Falcon tube (50 mL,) as provided, was treated with 1 mL 50% MeOH-Water. Vortexed thoroughly, sonicated (bath) for 30 min, centrifuged (3000 RPM, 15 min, 4°C). The supernatant was recovered into Glass vial (2 mL) and stored at −20°C overnight. The vials were centrifuged again centrifuged (3000 RPM, 15 min, 4°C). From the supernatant, 0.2 mL was taken out for GC/MS derivatization. From the supernatant, 0.04 mL aliquot were taken from each sample to generate pooled QC samples (7 x 0.2 mL).

#### Soil

Portions of submitted soil samples were frozen (−80°C) and lyophilized. The lyophilized soil samples were then pulverized using a bullet blender. ~ 150 mg of powdered soil samples were taken in Eppendorf tubes and treated with 1 mL water. Vortexed thoroughly, sonicated (bath) for 30 min, centrifuged (10000 RPM, 15 min, 4°C). From the supernatant, 0.75 mL taken out into new Eppendorf. Vortexed thoroughly, sonicated (bath) for 30 min, centrifuged (10000 RPM, 15 min, 4°C), frozen, and lyophilized. To these lyophilized samples, 0.75 mL 80% MeOH-Water with 0.1% formic acid was added. Vortexed thoroughly, sonicated (bath) for 10 min, centrifuged (10000 RPM, 15 min, 4°C). Stored at −20C for 2 h and centrifuged again (10000 RPM, 15 min, 4°C). From the supernatant, 0.5 mL for GC. From the supernatant, 0.1 mL aliquot were taken from each sample to generate pooled QC samples (14 x 0.5 mL).

#### Derivatization

All samples were dried under N2-stream. The dried samples were treated with 0.05 mL of methoxyamine hydrochloride in pyridine (25 mg/mL), vortexed and heated at 60°C for 45 min. The vials were then sonicated for 10 min and incubated again for another 45 min. The samples were then centrifuged at 2000 RPM for 5 min, treated with MSTFA+1%TMCS (0.05 mL), vortexed thoroughly and incubated at 60°C for 35 min. The samples were then put into inserts and analyzed by GC/MS.

#### GC-MS Data Acquisition

Metabolites were detected using a Trace 1310 GC coupled to a Thermo ISQ mass spectrometer. Samples (1 µL) were injected at a 10:1 split ratio to a 30 m TG-5MS column (Thermo Scientific, 0.25 mm i.d., 0.25 μm film thickness) with a 1.2 mL/min helium gas flow rate. GC inlet was held at 285°C. The oven program starts at 80°C for 30 s, followed by a ramp of 15°C/min to 330°C, and an 8 min hold. Masses between 50-650 m/z are scanned at 5 scans/sec under electron impact ionization. Transfer line and ion source are held at 300 and 260°C, respectively. QC samples were injected after every 6 experimental samples

#### Data analysis

Peak detection, alignment, and peak filling was performed on .cdf converted files using XCMS in R version 4.2.2. Additional feature clustering was performed using RAMClustR version 1.2.4 in was used to normalize, filter, and group features into spectra.XCMS (Smith 2006)(Tautenhahn 2008) output data was transferred to a ramclustR object using the rc.get.xcms.data function. Feature data was extracted using the xcms featureValues function. Features which failed to demonstrate signal intensity of at least 1.5 fold greater in QC samples than in blanks were removed from the feature dataset. 718 of 3117 features were removed. Features with missing values were replaced with small values simulating noise. The absolute value was used as the filled value to ensure that only non-negative values carried forward. Zero values were treated as missing values. Variance in quality control samples was described using the rc.qc function within ramclustR. Summary statistics are provided including the relative standard deviation of QC samples to all samples in PCA space, as well as the relative standard deviation of each feature/compound in QC samples, plotted as a histogram. Features were normalized by linearly regressing run order versus qc feature intensities to account for instrument signal intensity drift. Only features with a regression pvalue less than 0.05 and an r-squared greater than 0.1 were corrected. Of 2399 features, 66 were corrected for run order effects. Features were clustered using the ramclustR algorithm (Broeckling 2014). Parameter settings were as follows: st = 3, sr = 0.5, maxt = 300, deepSplit = FALSE, hmax = 0.5, minModuleSize = 2, and cor.method = pearson.

## Notes

### Competing Interest Statement

The authors have declared no competing interest.

### Summary of Updates

This version of the manuscript has been revised to update the following: author name corrected, a mistype on line 355 was corrected.

